# Local cryptic diversity in salinity adaptation mechanisms in a wild outcrossing *Brassica*

**DOI:** 10.1101/2024.04.18.590122

**Authors:** Silvia Busoms, Ana C. da Silva, Glòria Escolà, Raziyeh Abdilzadeh, Emma Curran, Anita Bollmann-Giolai, Sian Bray, Michael Wilson, Charlotte Poschenrieder, Levi Yant

## Abstract

It is generally assumed that populations of the same species should evolve shared mechanisms of adaptation to common stressors due to evolutionary constraint. Here, we describe a novel system of within-species local adaptation to coastal habitats, *Brassica fruticulosa,* and detail surprising mechanistic variability in adaptive responses to extreme salinity. These radically different adaptive responses in neighbouring populations are evidenced by transcriptomes, diverse physiological outputs, and completely distinct genomic selective landscapes. In response to high salinity Northern Catalonian populations restrict root-to-shoot Na^+^ transport, favouring K^+^ uptake. Contrastingly, Central Catalonian populations accumulate Na^+^ in leaves and compensate for the osmotic imbalance with compatible solutes such as proline and elevated Ca^2+^. Despite contrasting responses, both metapopulations were salinity tolerant relative to all inland accessions. To characterise the genomic basis of these two divergent adaptive strategies in an otherwise non-saline-tolerant endemic, we generate a long-read-based genome and population sequencing of 18 populations (9 inland, 9 coastal) across the *B. fruticulosa* species range. Results of genomic and transcriptomic approaches confirm the physiological observations of completely distinct underlying mechanisms of adaptation to extreme salinity and reveal potential genetic targets of these two recently evolved salinity adaptations. We therefore provide a new model of within-species salinity adaptation and reveal cryptic variation in neighbouring plant populations in the mechanisms of adaptation to an important natural stressor highly relevant to agriculture.

**Significance:** It’s usually expected that closely related populations of a given species should adapt to the same environmental stressor in the same way due to genetic or physiological constraints. However, this is not commonly tested due to practical constraints. Here we show that, even at the level of neighbouring populations, contrasting adaptive mechanisms control adaptive responses to extreme coastal salinity in a new plant model, *Brassica fruticulosa*, a close wild relative of many crops of worldwide importance. This indicates multiple options for engineering an agriculturally crucial adaptation: soil salinization. These results will be of great interest to not only those studying fundamental mechanisms of adaptation, but also resilience improvement in Brassica species.

## Introduction

It is generally expected that populations belonging to the same species should share specific evolved mechanisms of adaptation to the same stressors, whether these stressors are simple or complex. However, this expectation has not been sufficiently tested due to constraints on study systems, sampling, resolution, and scale. We thus lack an understanding of how often this assumed generality of species-wide adaptation mechanisms is violated in favour of diverse adaptations to uniform challenges within species. Here we test this expectation by taking a ‘hyper-local’ approach in the study of plant adaptation to coastal stressors, focussing on adaptation to high coastal salinity in a strip of coastline in Catalunya, Northern Spain. Previous work on local adaptation of *Arabidopsis thaliana* in this region detailed geographically and temporally fine-scale adaptive variation in fitness-related traits across environmental salinity gradients, even at the scale of a few (3-5) kilometres [1; 2]. This region is characterised by a dramatic positive gradient of soil salinity from inland to the coast, shaping plant species communities and driving the evolution of salinity tolerance mechanisms at the local population-(deme-) level [3]. Plant evolutionary responses to these conditions have been observed even in the selfer *A. thaliana* at fine (3-5km) scale, resulting in functionally adaptive variation [4]. Functional confirmation of this is evidenced by selective sweep of a hypomorphic ion transporter HKT1;1, which modulates Na^+^ leaf concentrations in response to rapid (monthly) temporal and spatial variation in salinity and rainfall [1].

Unfortunately, work in *A. thaliana* has two major limitations: first, due to its overwhelmingly selfing reproductive mode, relative to its outcrossing relatives *A. thaliana* has tenfold lower genetic diversity and high rates of spontaneous, population-specific mutations [5]. This low diversity also has important consequences in respect to increased homozygosity and effective population size resulting in genetic drift, reduced effective recombination rates, genomic background effects, and the fixation of maladaptive alleles (reviewed in [6]). Secondly, *Arabidopsis* is substantially divergent from important Brassica crops, limiting the translational potential of discoveries in this otherwise convenient lab model. Wild outcrossing plants, on the other hand, harbour much higher levels of genetic diversity, directly facilitating studies of adaptation [7]. Motivated by these considerations, we searched for wild Brassicaceae species with contrasting, recently-evolved (within-species) phenotypes in complex coastal adaptations, focussing specifically on salinity tolerance. This resulted here in the identification of a new model for local adaptation to coastal salinity, *Brassica fruticulosa*, and allows us to test hypotheses regarding the scale of local adaptation to high coastal salinity.

The genus *Brassica* belongs to the Brassicaceae (mustard) family and contains nearly 100 species, many of which are grown globally as vegetables like cabbage, broccoli, kale, and radish, as mustards, as oil crops (placing 3^rd^ after palm and soy) and as fodder for animal feed [8]. Brassicas are widely proficient at adapting to new habitats due to recent and recurrent polyploidy events, hybridisation, and plastic genomes. These characteristics also make them great targets for genetic manipulation to further enhance resilience [9].

Here, we first perform a large-scale, genus-wide natural variation survey of diverse, wild outcrossing Brassicas in coastal northeast Spain, eventually testing six candidate species for within-species adaptation to high salinity. From these, we identify and develop one particularly promising novel model of within-species variation in adaptation to extreme salinity and complex coastal stressors, *Brassica fruticulosa.* First described in 1792 by Cirillo [10], *B. fruticulosa* has not yet been recognized as harbouring population-specific salinity adaptation. This has been a missed opportunity, as *B. fruticulosa* is closely related to *Brassica rapa* [11; 12] and shares many affinities with this global crop. We then assemble an Oxford Nanopore-based *B. fruticulosa* genome, sequence 90 individuals from 18 populations (9 coastal, 9 inland) contrasting in salinity and soil parameters defined by ionome levels in leaves and soil in the root space of every individually sequenced wild plant. Using transcriptome data of leaves and roots, we reveal divergent adaptive strategies in response to high salinity in neighbouring plant populations. We then perform common garden, physiological, and ion homeostasis experiments to detail these differing strategies that evolved in closely neighbouring adapted plant populations. Finally, we perform environmental association analysis (with soil ionome as phenotype) and genome scans by ecotype to seek a genomic basis of divergent adaptation strategies to salinity tolerance in neighbouring *B. fruticulosa* populations. Taken together, these experiments reveal surprisingly contrasting adaptive responses to extreme salinity, at the local scale, differing mechanistically at the scale of kilometres.

## Results and Discussion

### A region-exhaustive screen for fine-scale salinity adaptation in Catalonia

Situated in northeast Spain, Catalonia harbours a diverse flora ranging across a positive gradient of soil salinity from inland to the coast. This salinity gradient acts as a selective agent upon coastal plants to develop mechanisms of adaptation to high coastal salinity [3], and is accompanied by other local stressors (such as impoverished soils, alkalinity or drought), which are understood in good resolution [13; 14; 15]. These selective agents, combined with the fact that the Iberian region has served as an important glacial refugium for many species, have resulted in dramatic local plant diversification. For example, in the most well-studied system in terms of genome resources, Iberian *A. thaliana* harbours the largest genomic diversity of the entire species distribution [16] and has shown significant adaptive variation in fitness-related traits across environmental gradients [e.g. 17; 18].

We thus performed a search for all wild Brassicaceae populations in the region with contrasting interspecies phenotypes in salinity tolerance. We first identified populations from 13 wild Brassica species and georeferenced these from available databases (methods). Six of these 13 species were found in more than 2 coastal and 2 inland sites. Seeds from these species (*B. fruticulosa*, *Lepidium graminifolium*, *Lobularia maritima*, *Diplotaxis tenuifolia, Diplotaxis erucoides,* and *Diplotaxis muralis*) were collected (Dataset S1) and progeny were tested for salinity tolerance. Salinity tolerance in *L. maritima*, *D. tenuifolia* and *D. muralis* was uniformly high in both coastal and inland ecotypes, with plants completing their life cycle under continuous exposure to 300 mM of NaCl regardless of their coastal or inland origin. Coastal and inland *D. erucoides* and *L. graminifolium* plants exhibited equally higher sensitivity, while still enduring 200 mM NaCl for 2-3 weeks but not reproducing after longer exposures (Dataset S1). Of all the Brassicaceae species recorded in the region and identified both inland and coastally, only *B. fruticulosa* exhibited a within-species contrast in the ability to survive in high soil salinity (Fig. 1A).

**Figure 1.**
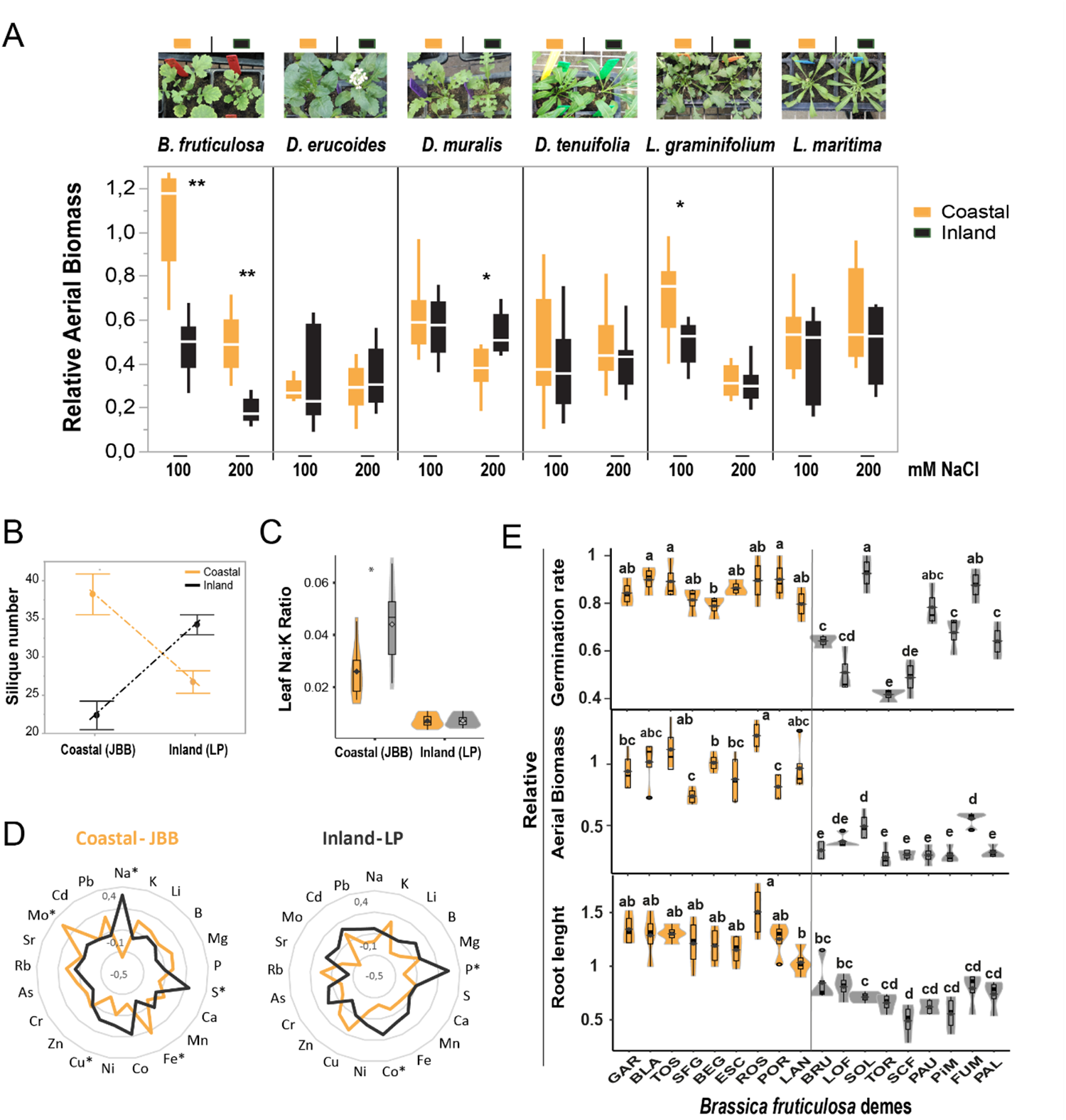
*Brassica fruticulosa* exhibits marked within-species variation in performance under salinity relative to other local Brassicaceae species in a region-exhaustive survey. (**A**) Relative aerial biomass (DW Treatment / DW Control) of *Brassica fruticulosa*, *Diplotaxis erucoides, Diplotaxis muralis*, *Diplotaxis tenuifolia, Lepidium graminifolium*, and *Lobularia maritima* plants from coastal (orange) and inland (black) populations cultivated in common soil irrigated with 100 mM NaCl or with 200 mM NaCl for one week. Asterisks indicate significant differences (ANOVA, *p<0.1, **p<0.05) in aerial biomass between origin under these saline conditions. Pictures representative of coastal and inland 21-day-old plants treated with 100 mM NaCl are above. (**B**) Fitness (silique number as proxy), (**C**) leaf Na:K ratio, and (**D**) leaf ionome profile of *B. fruticulosa* plants from coastal (orange) and inland (black) populations cultivated in a coastal field common garden (JBB) and an inland field common garden (LP). (**E**) Relative (Treatment/Control) germination rate, aerial biomass, and root length of *B. fruticulosa* populations sown in MS plates (0 or 75 mM NaCl). Seedling germinated without NaCl were transferred to hydroponic conditions and treated with 0 mM (Control) or 150 mM NaCl (Treatment) for 10 days. Letters indicate significant differences between populations (Tukeýs HDS test).

### Local adaptation and intraspecific variation for salinity tolerance in *Brassica fruticulosa*

Having discovered marked salinity tolerance of particular *B. fruticulosa* populations under controlled laboratory conditions, we tested for local adaption to inland and/or coastal sites. We performed a reciprocal transplant experiment in two representative local field sites (JBB-coastal and LP-inland), which had been established for similarly testing local adaptation in wild *A. thaliana* [3; 18]. As a proxy for the fitness of each plant, we counted the number of siliques produced. Additionally, thirty days after germination one leaf per plant was harvested for ionomic analysis. In across-site comparisons, both coastal and inland plants performed better in their home environment (Fig. 1B). In within-site comparisons, coastal plants outperformed inland plants when both were grown together in the coastal environment, whereas inland plants displayed a significant fitness advantage compared with coastal plants when both were grown inland (Fig. 1B), indicating that both the coastal and inland populations are locally adapted.

Having established this local adaptation of coastal and inland populations, and in particular the relative salinity tolerance of coastal populations (Fig. 1A), we then focussed on possible mechanisms underlying the high salinity tolerance of the coastal populations. This increased tolerance may be caused by either Na^+^ exclusion or the accumulation of Na^+^ and internal tissue tolerance mechanisms [4; 19]. Whole leaf ionome analysis revealed that *B. fruticulosa* plants from coastal and inland populations did not differ in their accumulation of Na^+^ or K^+^ under inland conditions. However, in coastal common gardens, inland plants accumulated more Na^+^ than coastal plants, resulting in a significant increase in the Na:K ratio (Fig. 1C&D, Dataset S3). These results indicate that coastal populations restrict leaf Na^+^ accumulation, maintaining a significantly lower Na:K ratio under coastal conditions. Conversely, at the inland site, most of the essential nutrients, such as P, were higher in the *B. fruticulosa* inland populations (Fig. 1D), reflecting the better growth and nutritional status of inland plants in their local origin.

To explicitly test for intraspecific salinity tolerance of all *B. fruticulosa* populations, a hydroponic experiment was conducted to ensure uniform NaCl treatment. Thirty seeds from each population were placed in square plates with ½ MS medium (pH 6) or ½ MS medium with 75 mM of NaCl (pH 6). One-week-old seedlings from the control plates were transferred to a hydroponic system filled with 0.5-strength Hoagland solution (pH 6). After a week, salinity was gradually increased (until a maximum of 150 mM NaCl, pH 6) in the group of treated plants. More than 75% of seeds from coastal populations germinated under saline conditions.

Contrastingly, 6 of the 9 inland populations had a relative germination rate below 70% (Fig. 1E). Some inland populations (SOL, PAU and FUM) could germinate well in 75 mM NaCl, but already at the seedling stage they suffered a significant reduction of growth and root development when exposed to salinity (150 mM of NaCl; Fig. 1E). All *B. fruticulosa* coastal populations outperformed the inland populations in terms of aerial biomass (Fig. 1E), explicitly identifying salinity tolerance in these populations.

### Long read-based *B. fruticulosa* genome assembly and annotation

As a first step in determining the genomic basis of the observed variation in salinity tolerance in *B. fruticulosa*, we generated a reference genome of an individual from a particularly salinity-tolerant population, TOS (Tossa de Mar, Catalonia, Spain), using long-read Oxford Nanopore Technology (ONT). Libraries were sequenced on the PromethION platform and base-called with the high accuracy model in Guppy (version 6). For assembly, we used 1,806,322 ONT reads with a read length N50 of 31 kb (21 total GB ONT data). This represents 38x coverage of the estimated 557 megabase genome (the kmer-based size estimate by FindGSE [20]). Using Flye (version 2.9), we first produced a pseudohaploid assembly of 505 mb. We then polished the assembly with Medaka [21] and Pilon [22], incorporating paired-end Illumina data, and then purged duplicate uncollapsed haplotypes with purge_dups [23]. We removed contaminating sequence by using the BlobTools pipeline [24] using coverage, Diamond-blast, and GC content information, with results shown in Dataset S5. This resulted in a final purged assembly of 351 mb with an N50 of 470 kb (Fig. 2A). Given a repetitive ratio of 0.56 (312 mb), we expect this assembly to be complete for the non-repetitive gene space. This was confirmed through benchmarking analysis using BUSCO [25] against brassicales_odb10. This showed high completeness (97.1% complete BUSCOs; Fig. 2A). Finally, gene models were predicted by an evidence-guided annotation approach incorporating RNA-Seq data and cross-species protein alignments in BRAKER2 [26].

**Figure 2.**
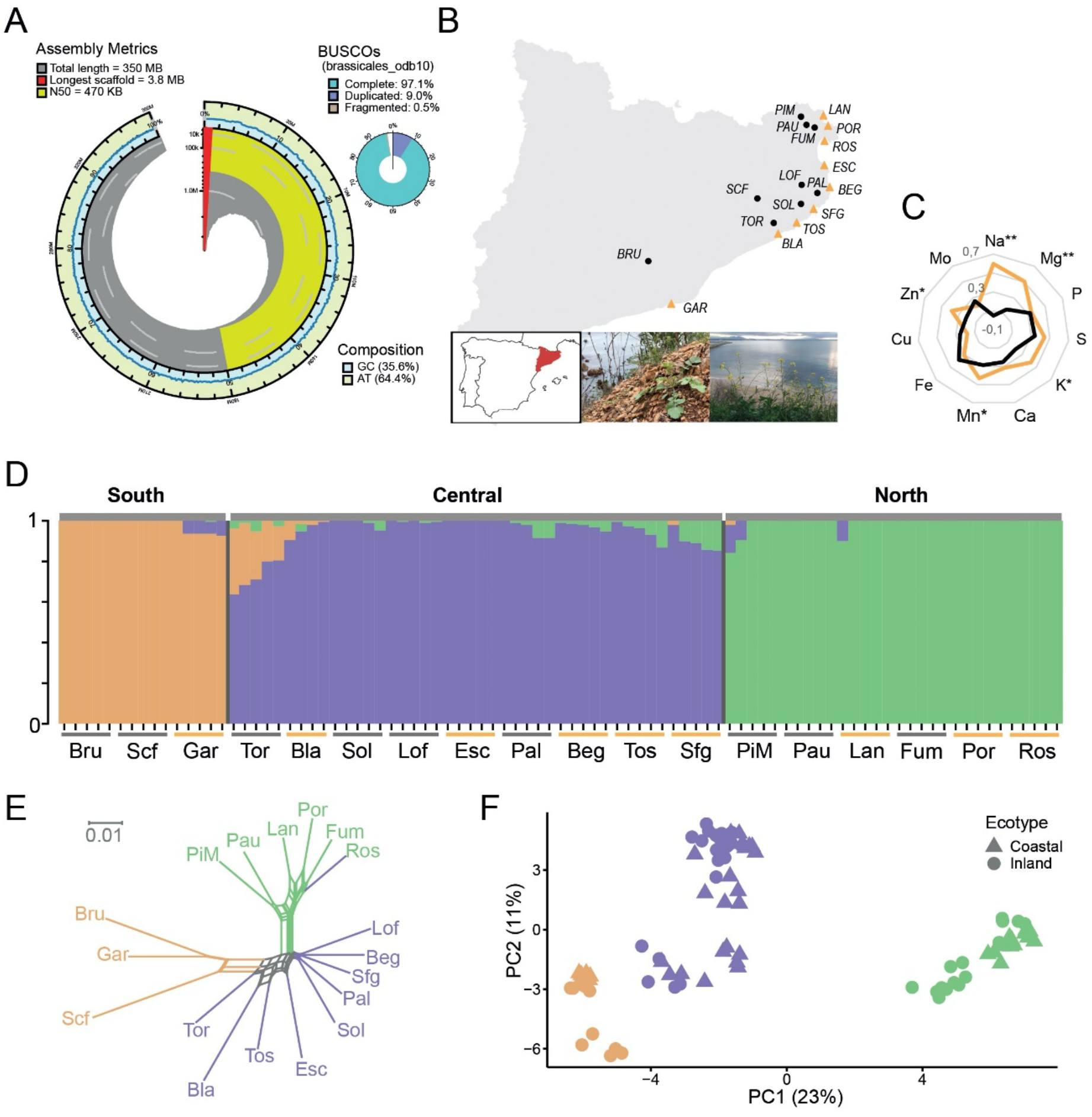
Long-read genome assembly, population resequencing, and genetic structure of *B. fruticulosa*. **(A)** Genome assembly and gene content metrics of Oxford Nanopore-based *B. fruticulosa* assembly. **(B)** Sampling map of Catalonia, Northern Spain, of the 18 populations sequenced. Inset: image of saline-tolerant coastal *B. fruticulosa.* **(C)** Radar plot representing the concentration of 11 elements in the soil of inland (black line) and coastal (orange line) populations. Asterisks indicate significant differences between origin (ANOVA, *p<0.05, **p<0.001)**. (D)** fastSTRUCTURE analysis (k=3, min MAF 2.5%) of the 89 individuals analysed, with clusters corresponding tightly to geographic origin, but not to salinity tolerance or coastal/inland status. **(E)** Nei’s genetic distances between populations visualized in SplitsTree. **(F)** PCA of all individuals. Coastal populations are represented by triangles and inland populations by circles. Northern populations are coloured in green, central in purple, and southern in orange.

### Population sequencing and genetic structure analysis

To assess population- and individual-level demographic and selective landscapes, we then sampled 18 populations (Fig. 2B) for genome sequencing across the reported range of *B. fruticulosa*. We Illumina sequenced a total of 90 individuals, selecting 5 individuals from each of the 18 populations (average depth per individual = 13.2x, post-quality filtering; 10,254,275 variable SNPs on average; Dataset S5). One sample, Bla5, was excluded, due to suspicion of sample contamination.

To assess genetic structure we performed fastSTRUCTURE [27] and NGSadmix [28] (Fig. 2C) on our 89 individuals (30,272 LD-pruned, biallelic 4-fold-degenerate SNPs; max 20% missing data; min MAF = 2.5%). K=3 maximised marginal likelihood, consistent also with Discriminant Analysis of Principal Components (DAPC) analysis of the Bayesian Information Criterion in *adgenet* [29]. The K=3 result reflects a northern group (including 3 coastal and 3 inland populations), a central group (including 5 coastal and 4 inland populations), and a very small southern group (with only 2 inland and 1 coastal population) that was excluded in the following analysis. This grouping partitioned all samples by geographic region of origin (Fig. 2B&D-F) and not by salinity tolerance status or local soil characteristics (Fig. 2C). Both Nei’s simple genetic distances visualised in SplitsTree (Fig. 2E) and PCA (Fig. 2F) confirmed that geographic origin completely dominates over saline/non-saline ecotype.

### Highly divergent functional diversity in salinity responses at the local level

Given our recognition of two larger genetic clusters, consistent with local origin (here termed the ‘north’ and ‘central’ metapopulations) we performed a salinity challenge experiment to test the transcriptome response of each population. We chose the most sharply contrasting populations from each major genetic cluster (north and central) from the hydroponic salinity challenge experiment above and then subjected them to salinity challenge (0 mM or 150 mM NaCl) for 10 days. Leaves and roots of 6 plants per population (48 samples total: 2 tissues of 24 plants from 2 salt-tolerant and 2 salt-sensitive populations, 12 plants under control and 12 plants under salt stress conditions) were sequenced for transcriptome analysis.

PCA of the overall transcriptome response revealed a surprising marked differential effect of genetic cluster on salinity response profiles, specifically in leaves. While under control conditions, all samples cluster together regardless of population origin or salt sensitivity (black shapes in Fig. 3A), upon salt treatment (yellow shapes in Fig. 3A), plants from the ‘north populations’ (circled in green) and plants from the ‘central populations’ (circled in purple) show divergent transcriptomic responses along both PC1 and PC2. This points to a specifically salinity-related divergence in genomic responses between these two salt resistant metapopulations at the transcriptomic level. This also provided a first hint at divergent salinity tolerance mechanisms between the north and central metapopulations, despite close proximity in nature and very low genetic divergence between the north and central coastal populations (pairwise Fst = 0.13).

**Figure 3.**
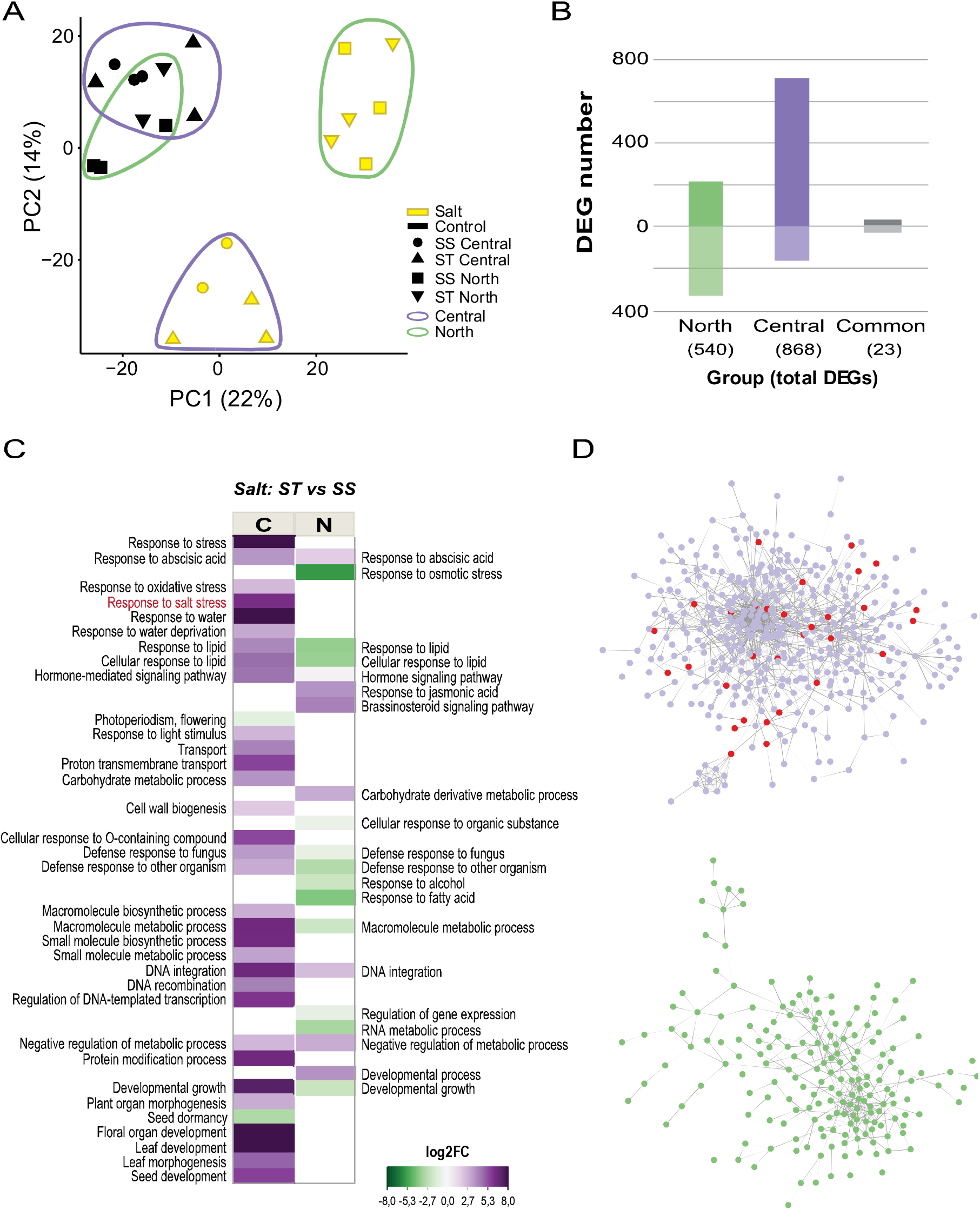
Differential transcriptome responses to salinity challenge between *Brassica fruticulosa* north and central metapopulations. **(A)** Principal component analysis (PCA) of leaf transcriptome profiles of 12 salt-tolerant (triangles up/down) and 12 salt-sensitives (circle/square) *B. fruticulosa* individuals from north (green outline) and central populations (purple outline) treated with 0 mM NaCl (control, black) or 150 mM NaCl (salt, yellow) for 10 days. Root data are given in Figure S1. **(B)** Number of differentially expressed genes (DEGs) in pairwise comparations Salt Tolerant (ST) *vs* Salt Sensitive (SS) under salt stress of north populations (green bars), central populations (purple bars), and shared in both origins (grey bars). **(C)** Significantly enriched gene ontology (GO) terms (biological process category, using a Fisher’s exact test with a p-value ≥ 0.05) associated with DEGs from salt tolerant vs salt sensitive north comparison (right panel), and from salt tolerant vs salt sensitive central populations comparison (left panel). Scale colour indicates the L2FC mean of the genes included in each GO term. (**D**) Probabilistic functional gene networks of DEGs from the central comparison (purple circles) and from the north (green circles). ‘Response to salt stresś subcommunity is indicated with red circles.

Considering this clearly contrasting salinity response in each group revealed by PCA, we analysed transcriptome responses separately for north or central populations, hypothesizing that salt-tolerant plants might be using different mechanisms to cope with salinity stress. Doings so we found that the quantity and identity of differential expressed genes (DEGs) differed radically between the north and central metapopulations. The analysis of salt tolerant *vs* salt sensitive *B. fruticulosa* plants from the north populations (green in Fig. 2D-F, Fig. 3B) gave 540 DEGs (L2FC > |2|, p-adj < 0.05) activated in leaves only under salt stress conditions, with 213 up- and 327 down-regulated. In the same analysis performed with the central populations (purple in Fig. 2D-F, Fig. 3B) we obtained a total of 868 DEGs, 708 up- and 160 down-regulated under salinity (Dataset S6). Many fewer DEGs shared the same expression pattern in both analyses (only 23 DEGs; corresponding to 2.6% of DEGs in central and 4.3% of DEGs in northern), which constitutes a very small minority of the DEGs in each cluster (Fig. 3B). This confirms the observation of contrasting global transcriptional salinity stress responses as a function of genetic cluster by PCA (Fig. 2C).

Both the greater quantity of upregulated genes detected in the central cluster, together with the gene ontology (GO) terms enriched (Fig. 3C, Fig. S2), indicate that salinity stress activates several biological responses to cope with abiotic stresses specifically in the central populations: transport control, hormone regulation, decrease of oxidative stress, and responses to water deprivation. The transcriptomic analysis (Dataset S6) provides highly valuable information guiding future characterization of the genes potentially involved in salt tolerance based on internal detoxification in the *B. fruticulosa* populations from the central cluster. Here we point out only a few examples of highly upregulated genes of interest, based on homology to *A. thaliana* and other brassica relatives where these have been studied. Involved in the alleviation of oxidative stress, among others, we find glutathione transferases (GSTU3, 6, 11, 13, and 25) and alternative oxidases AOX1A and AOX1B (22- and 3-fold, respectively); in transport the vesicle transporter SEC1A (20-fold), the fluoride exporter FEX (6-fold), the Mg transporter AT2G04305.1 (7-fold), and the vacuolar transporter ABCG19 (9-fold) are notable. In stomatal regulation and drought and salt responses DUF810 (20-fold) and YL1/BPG2 (11-fold), both involved in salt stress through ABI14 regulation, stand out.

In stark contrast, most of these stress response terms are not enriched, or are downregulated, in the north populations under salinity challenge (Fig. 3C&D, Fig. S2). This indicates that, relative to salt tolerant central plants, salt tolerant northern plants are not sensing stress caused by Na^+^ toxicity. However, secondary metabolism and hormonal signalling pathways are clearly highly activated through upregulation of ERF transcription factors (ERF/AP2, 6-fold) and other genes involved in ABA (EDL3, 10-fold) and Brassinosteroid (BR6OX2, 4-fold) signalling. Other examples of strongly upregulated genes in the tolerant northern cluster are, among others, sterol isomerase (AT1G05440, 9-fold); boron efflux transporter BOR4 (7-fold); SNF1, involved in osmotic and ionic stress response (7-fold); or OXR1, an oxidative stress tolerance gene (7-fold) (Dataset S6).

To better resolve potential mechanisms used by salt tolerant northern and central plants, salt stress-diagnostic physiological parameters were evaluated in the same individuals used for the transcriptome analysis. Overall, salt tolerant plants maintained higher osmotic potential and exhibited greater growth than salt sensitive plants, independently of cluster origin (Fig. 4A-C). However, salt tolerant northern plants were much more efficient at restricting Na^+^ translocation and favouring K^+^ uptake than salt tolerant central plants (Fig. 4E&F). Indeed, when we tested expression levels of well-studied orthologs involved in Na^+^ and K^+^ transport, we observed that HKT1 was significantly upregulated especially in the roots of salt tolerant plants from north populations (Fig. 4H). It has been seen in several species that high expression levels of HKT transporters in the root leads to lower shoot Na^+^ concentrations due to the role of HKT in retrieval of Na^+^ from the transpiration stream [30]. Also, a repressor of HKT1, PP2C49, was downregulated (Fig. 4H), probably enhancing shoot Na^+^ extrusion, as shown in *A. thaliana pp2C49* mutants [31]. In addition, high salinity usually causes the downregulation of HAK/KUP/KT family members such as HAK5 in roots [32]. However, in salt tolerant northern *B. fruticulosa* plants the expression of HAK5 in roots was not altered and the gene was upregulated in the leaf tissue (Fig. 4H), suggesting that K^+^ uptake and translocation may be promoted preventing Na^+^ accumulation.

**Figure 4.**
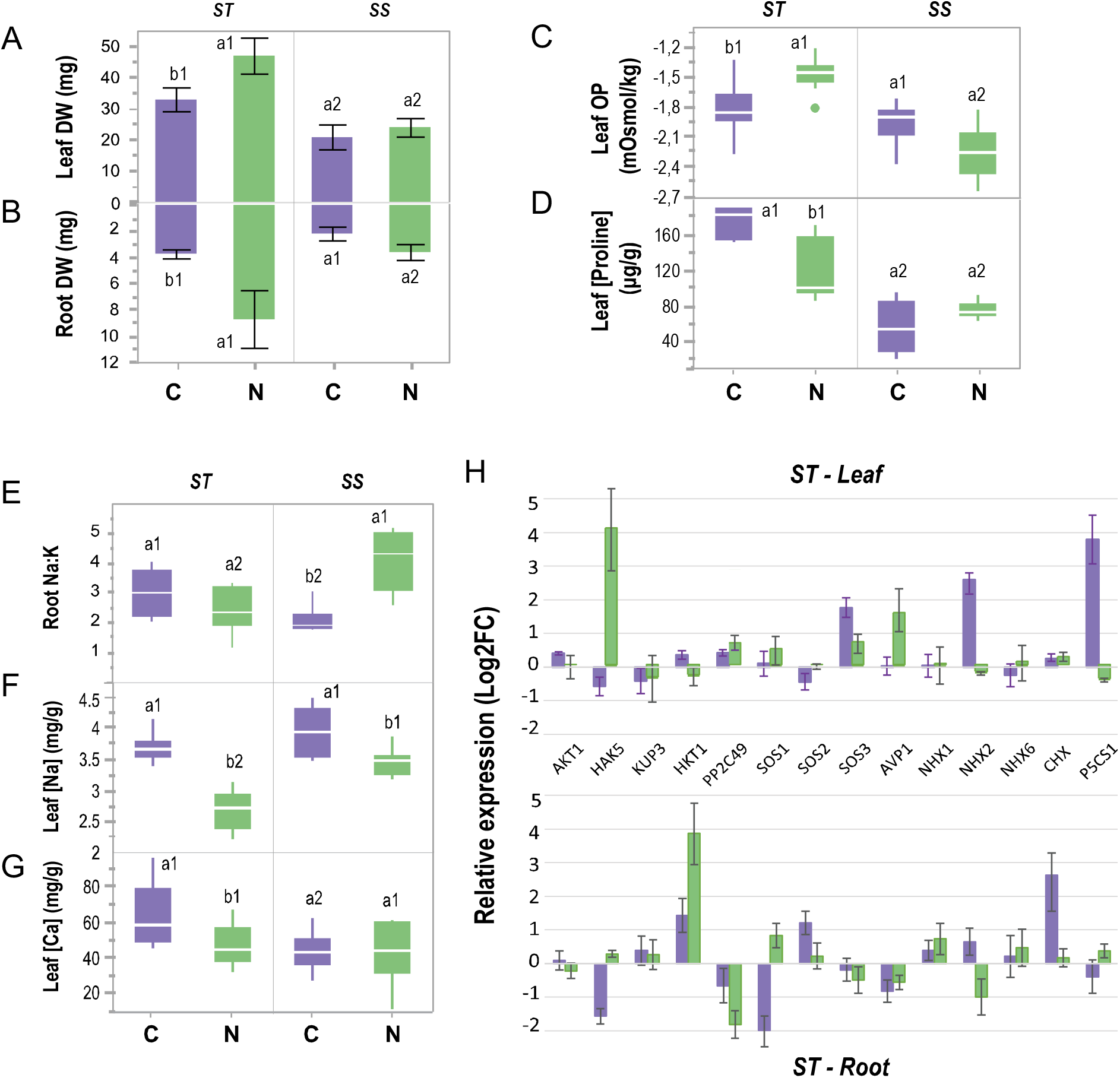
Disjunct adaptive phenotypic and genotypic responses of north and central *B. fruticulosa* populations. **(A)** Shoot and **(B)** root biomass (DW, g); **(C)** leaf osmotic potential (mOsmol/Kg), **(D)** leaf [Na^+^](mg/g), **(E)** leaf [Ca^2+^](mg/g DW), **(F)** leaf [proline] (microgram/g FW), and **(G)** root Na:K ratio of salt sensitive (SS) and salt tolerant (ST) *B. fruticulosa* plants from north (green) and central (purple) populations cultivated hydroponically and treated with 0 mM NaCl (control) or 150 mM NaCl (salt) for 10 days. **(H)** Root and leaf relative expression (Log2FC salt *vs* control) of 14 orthologs involved in Na^+^ and K^+^ transport or salinity responses in salt tolerant *B. fruticulosa* plants. Letters indicate significant differences between geographic origin and numbers indicate significant differences between salinity tolerance phenotype (ANOVA, p<0.05).

Contrastingly, salt tolerant central plants accumulated higher Na^+^ levels in the leaves, but the osmotic imbalance was compensated with a higher production of compatible solutes such proline (Fig. 4D). Influx of Na^+^ into the root system is counteracted by efflux of Na^+^ to the rhizosphere, an active process that occurs in antiport with H^+^. This process involves the *Salt Overly Sensitive* (SOS) pathway in many plant species, in which the calcium-binding protein SOS3 recruits the Ser/Thr protein kinase SOS2 to the plasma membrane where it activates the Na^+^/H^+^ antiporter SOS1 by phosphorylation [33]. SOS1 was downregulated in salt tolerant central roots (Fig. 4H), showing the inverse relationship between SOS1 expression in the root and total plant Na^+^ accumulation also found in other brassicas [34; 35]. Thus, the Na^+^ efflux in salt tolerant central plants is low but they must have the ability to tolerate Na^+^ inside their tissues. Vacuolar storage of Na^+^ is mediated by vacuolar Na^+^/H^+^ antiporters (NHX clades) and the electrochemical potential is provided by vacuolar H^+^-pyrophosphatases (e.g. AVP1) and vacuolar H^+^-ATPase [36]. Indeed, *NHX2* was significantly upregulated in the leaves of salt tolerant central plants (Fig. 4H) and could be enhancing Na^+^ vacuolar storage. Nonetheless, we also detected an overexpression of a cation/H^+^ exchanger (CHX), a putative K^+^, Na^+^/H^+^ antiporter, in the roots of salt tolerant central plants (Fig.4H). The CHX genes of *A. thaliana*, soybean, rice, and other plants have been identified as being involved in tolerance to salt stress thanks to their role in K^+^ and Na^+^ acquisition and homeostasis [37]. Recently, Guo et al. [38] showed that overexpression of a halophyte CHX gene (*KvCHX*) in *A. thaliana* seedlings enhanced their salinity tolerance by increasing K^+^/Na^+^ and proline levels.

Salt tolerant central plants also harboured elevated Ca^2+^ levels in leaves (Fig. 4G), suggesting that Ca^2+^ waves and root-to-shoot signalling are more active in these plants. Salt stress and changes in osmotic pressure are associated with the elevation of Ca^2+^ in the cytosol and the activation of ROS signalling. These signals induce adaptive processes to alleviate Na^+^ toxicity, including the maintenance of ion balance, the induction of phytohormone signalling, and increased synthesis of osmolytes and antioxidant enzymes [39, 40]. Indeed, the orthologue of P5CS1 (a synthase that catalyses the rate-limiting enzyme in the biosynthesis of proline), was highly expressed in salt tolerant central plants (Fig. 4H). P5CS1 appears to be involved in salt stress responses related to proline accumulation, including protection from reactive oxidative species [41], and may be crucial for *B. fruticulosa* salinity tolerance.

### Selection signatures associated with contrasting saline coastal adaptation

Having established that local adaptation to coastal environments is associated with two contrasting adaptive salinity tolerance mechanisms, we sought to further determine which alleles might underlie these phenotypes. Adaptation to coastal environments is complex and multidimensional, so we expect a diversity of selective pressures; further, given our differential salinity adaptation phenotypes between the north and central clusters, we expect particular regions to vary. We therefore applied a diversity of tests for selection. We first leveraged our population resequencing to search for footprints of selective sweep between coastal and inland populations. Given the extremely low within-group genetic divergence (mean saline/non-saline Fst = 0.016-0.019) and the lack of bottleneck (neutral Tajima’s D in all regions), this approach should avoid artefacts caused by bottlenecks or excess population divergence. For each cluster (north and central), we measured Hudson’s Fst in 1kb windows between saline and non-saline populations and from this extracted the 1% extreme outliers from the empirical distribution; these were reserved as our ‘Selective Sweep Candidates’ (‘Fst Outliers’, Fig. 5A and Dataset S8).

We then focussed on particular soil elements that significantly vary between coastal and inland sites (soil Na and Mg, Dataset S2), using these discriminant soil ion values as phenotypes in a targeted environmental association analysis (EAA). Here we inferred candidates directly associated with the chemical characteristics of each sequenced plant’s immediately proximal soil by performing EAAs using LFMM2 [42]. This analysis quantitatively determines the association between each soil element concentration and SNPs across the genome in all populations at the level of individual plants (north= 30; central=44). We identified 433 (north) and 56 (central) genes harbouring ≥1 SNP significantly associated with at least one distinctive soil parameter identified (‘EAA Outliers’, Dataset S9).

Finally, we performed a genome scan for regions of the genome that are outliers vis-à-vis background demographic structure with *PCAdapt* [43]. Such an approach provides an orthogonal test relative to the above, as it uses no priori information regarding group structure (as Fst scans do) or proposed selective agent/correlate (as EAAs do), instead identifying outlier regions based on a genome-wide Mahalanobis test statistic (‘PCAdapt Outliers’, Dataset S10).

**Figure 5.**
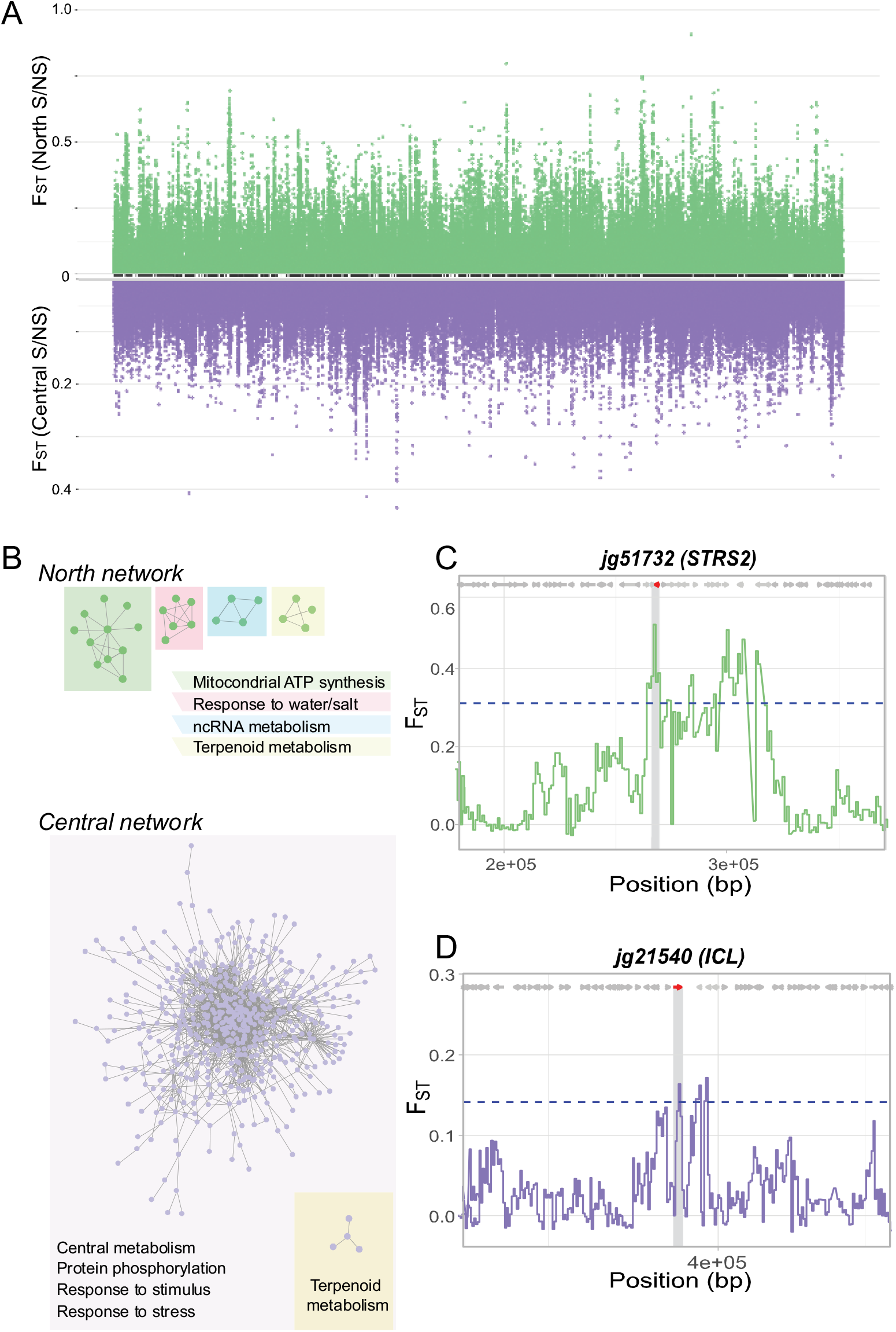
Completely divergent selection signatures in two adjacent saline-adapted *B. fruticulosa* metapopulations. **(A)** Contrasting patterns of ploidy-specific differentiation across the *B. fruticulosa* genome. Top Manhattan plot (green) shows F_ST_ values between coastal and inland populations in the north genetic cluster; bottom Manhattan plot (violet) shows F_ST_ values between coastal and inland populations in the central genetic cluster on an inverse y-axis in order to closely illustrate completely divergent selective sweep signals across the entire genome (x-axis). (**B**) Probabilistic functional gene networks of Fst 1% outliers from the North (green circles) and the Central (purple circles) clusters. Subcommunities are annotated according to significant enrichment in Gene Ontology terms for biological processes (adj-pval < 0.05, Fisher’s exact test). (**C-D**) Estimates of F_ST_ for selective sweep candidates from the north (**C**) and central (**D**) clusters. Solid lines give localised elevated F_ST,_ which is accompanied by low Tajima’s D driven sharply negative in the focal, coastalpopulation, a classic signal of selective sweep. All genes are F_ST_ 1% outliers related to salinity, jg51732 (AT5G08620) encodes a STRESS RESPONSE SUPPRESSOR 2 (STRS2) that binds to RNA and is involved in drought, salt and cold stress responses; jg21540 (AT3G21720) encodes a glyoxylate cycle enzyme isocitrate lyase (ICL) involved in salt tolerance.

Each of these approaches (transcriptome response, Fst outliers, EAA outliers, and PCAdapt outliers) yielded relevant candidates (Fig. 5; Datasets S6-10). But, as these approaches focus on orthogonal signals, there was limited concordance (Fig. S3). However, we do observe overlap at the level of GO enrichment within each genetic cluster. This was significantly greater overlap than expected by chance, as determined by permutation tests (p< 0.05, Dataset S11). Common relevant GO terms indicated by multiple approaches include ‘response to osmotic stress’ in the north cluster, and ‘response to water deprivation’ and ‘response to salt stress’ in the central cluster, both relevant to salinity adaptation (Fig. S3C-D, Dataset S11). Candidate genes contributing to these categories in our top candidate lists include the Plasma membrane Intrinsic Protein 1 (PIP1) and Stress Response Suppressor 2 (STRS2) for the north cluster and Tonoplast Intrinsic Protein 2 (TIP2), Salt Tolerance 32 (SAT32), Sugar Insensitive 8 (SIS8), UDP-Glycosyltransferase 79 (UGT79B2), and Isocitrate Lyase (ICL) for the central cluster. Membrane and tonoplast aquaporins such as PIP1 and TIP2 play crucial roles in mitigating salinity stress by maintaining water and ionic homeostasis and better cell membrane integrity [44]. Mutations in the DEAD-box RNA helicases STRS1 and STRS2 caused increased tolerance to salt and osmotic stress in *A. thaliana* by attenuating the expression of stress-responsive transcriptional activators [45]. Curiously, SAT32, associated with salt tolerance and ABA signaling in *A. thaliana*, has six copies and high structural variation in the halophytic *Eurtrema salsugineum* [46]. *A. thaliana* SIS8 is a MAPK that interacts with UDP-glucosyltransferase in the nucleus and negatively regulates salt tolerance [47] while overexpression of UGT79B2/B3 enhanced plant tolerance to salt stresses modulating anthocyanin accumulation [48]. In rice, ICL is regulated by a calcium sensor calmodulin (CaM)-encoding gene. The overexpression of both confers salinity tolerance in *A. thaliana* and rice by shifting energy metabolism from the TCA cycle to the glyoxylate cycle during salt stress [49].

### Evapotranspiration as a potential selective agent in a ’hyper-locaĺ scale

It was straightforward in *B.fruticulosa* to demonstrate local adaptation to coastal conditions as well as specific tolerance to elevated salinity. But we note also that adaptation to coastal environments is complex and multidimensional, with challenges to colonists not only from elevated salinity, but also various nutrient deficiencies and dehydration due to increased substrate porosity, wind exposure, sand-loaded sea spray, and high light intensity among other challenges [50]. Accordingly, signatures of selection may be expected to originate from stressors both common across coastal sites, but also distinctive to particular regions and even at the very local, microsite-scale (e.g. the scale of tens of meters; [51]). Thus, given the markedly contrasting genomic, physiological, and adaptive responses to the primary selective agent, salinity, we finally sought whether there may be subtle local differences in environmental challenge that may drive divergent responses to the ostensible primary agent of coastal salinity that were less obvious than the strong coastal salinity gradient across this region. Classic literature indicating a ‘hyper-local’ scale of plant adaptation further suggested fine-scale responses to stressors in adjacent plant populations e.g. on coastlines or mines [52]. These works found that especially in sedentary plants, selection can cause extremely localised patterns of micro-geographical adaptive variation despite a degree of migration; however, these processes depend on various ecological and genetic factors that are very challenging to evaluate, and unique for each species and habitat.

In our study region soil elemental composition does not vary between north and central coastal locations (Dataset S2); therefore, we looked more closely at climatic data, despite the highly local scale. While precipitation and daily temperatures did not differ between regions in the 14 years evaluated, evapotranspiration did (Fig. S4, Dataset S12). Evapotranspiration (ET0) is the sum of plant transpiration and soil evaporation and is affected by many environmental factors such as net radiation, air temperature, relative humidity, and especially wind speed [53]. Indeed, relative to the central region, tramontane wind is prevalent in the far north of Catalonia, and this explains why annual mean ET0 was significantly higher there, despite no substantial difference in precipitation or daily temperatures (Fig. S4, Dataset S12). An increased evaporative demand tends to increase transcriptional volume flow, leading to greater salt damage on plants [54]. Under these circumstances, we speculate that the observed Na^+^ exclusion strategy of the north cluster may be more effective. We speculate therefore that harsh coastal conditions aggravated by high ET0 are the primary agents of selection affecting *B. fruticulosa* northern region plants, resulting in selection on genes that confer advantage under drought and salinity stress. Contrastingly, in the central region, water shortage is less severe, and plants can cope with salinity with tissue tolerance and osmoprotection strategies (Figure 6).

**Figure 6.**
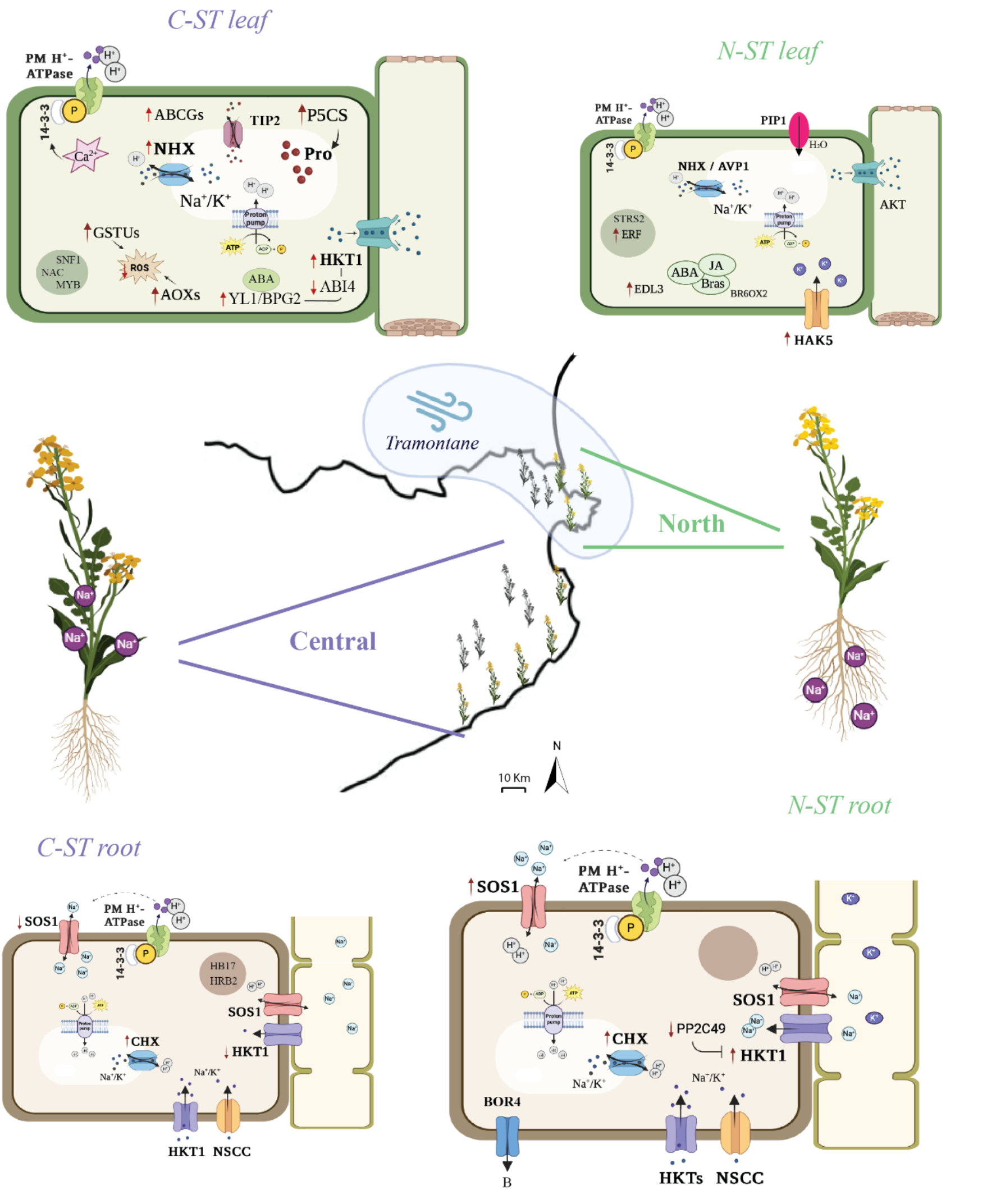
Overview of the contrasting salinity tolerance mechanisms of the North and Central *B. fruticulosa* coastal metapopulations. Gene symbols are shown in bold letters. Ion fluxes are indicated with black arrows. Gene activation/repression and molecule increase/decrease are indicated with red arrows. Star-framed symbols denote signalling pathways and hormone molecules are circled in green. ‘ST’=Salt tolerant.

## Conclusions

Here we set out to develop a novel model of within-species salinity tolerance in an outcrossing Brassicaceae, first by an exhaustive survey of 13 species in an ecologically well-understood region coastline in Northeast Spain. This search resulted in the discovery of two distinct evolved mechanisms of salinity tolerance in adjacent metapopulations of the same species, *Brassica fruticulosa* (Fig. 1). These metapopulations genetically cluster in 3 groups regardless of their coastal or inland origin and deploy contrasting mechanisms to mitigate salinity stress on the order of a dozen kilometres (Fig. 2). Relative to central salt tolerant plants (C-ST), northern salt tolerant plants (N-ST) excluded Na^+^ to a higher degree, restricting its translocation to the aerial tissues and favouring K^+^ uptake (Fig. 4 & 6). After 10 days under high salinity, root growth and synthesis of secondary metabolites were stimulated, key ion transporters such as SOS1 and HKTs were respectively down- and up-regulated in roots, and salt-responsive genes were not activated in leaves, suggesting that N-ST plants were not engaged in a salinity stress response (Fig. 3, 4, & 6). In contrast, C-ST did not restrict Na^+^ translocation, and accordingly instead activated the osmoprotectant machinery to compartmentalize Na^+^ in the vacuoles, thus reducing oxidative damage (Fig. 3, 4 & 6).

These divergent salinity responses were evidenced also in a variety of genome scans for the genomic basis of these contrasting responses (Fig. 5). We identify candidates for a mechanism of Na^+^ exclusion/extrusion and for a mechanism of effective Na^+^ compartmentalization, two strategies highly relevant for cultivars that need to adapt to the new environmental inclemencies. This work therefore establishes *B. fruticulosa*, a wild relative compatible with crops *Sinapis alba* and *Raphanus sativus*, as a promising source of wild desirable alleles, as well as these highly diverse natural populations as powerful new models for the study of adaptations to saline soils.

Our observed contrast in evolved ecophysiological salinity tolerance strategies between the north and central metapopulations is an interesting problem. These populations are situated on a coastal strip with a marked uniform salinity gradient, acting as obvious selective agents across the study region. However, this clearly divergent adaptive response of two adjacent populations required us to question whether other latent stressors may modify primary adaptive strategies. While soil salinity appears to be the principal agent driving coastal adaptation of *B. fruticulosa* populations, we speculate that other microclimate factors like variation in evapotranspiration rates may drive contrasting salinity tolerance strategies (Fig. 6). Future studies should work toward fully decoupling the complex selective forces responsible for these contrasting mechanisms.

## Materials and Methods

### Plant and soil materials

Seeds of wild Brassicaceae species were collected in Spring 2017 across Catalonia (map: https://www.google.com/maps/d/u/0/edit?mid=1PGHyv3p7AwtI57N6C_eC6ecbnj0&usp=sharing). Each population was classified depending on coastal (< 3 km from the sea) or inland origin. The 6 brassica species with 2 or more inland and coastal populations were selected for testing salinity tolerance (*Brassica fruticulosa* (BF), *Lepidium graminifolium* (LG), *Lobularia maritima* (LM), *Diplotaxis tenuifolia* (DT)*, Diplotaxis erucoides* (DE), and *Diplotaxis muralis* (DM)). Seeds were stored in cold and dry conditions until use.

Leaves of *Brassica fruticulosa* plants from 19 populations were collected and brought to the lab. Leaf samples from 6 individuals per population were dried for 48 hours at 60 °C and stored in a drier for the following ionomic analysis. Leaf samples from 5 individuals per population were frozen and kept at -20°C for subsequent DNA extraction and sequencing. Soil samples from near 3 *B. fruticulosa* plants separated at least 2 m were collected for each of the 19 populations, air-dried for 2 days, 2 mm sieved, and stored in cold/dry conditions for further soil ionomic analysis.

### Soil cultivation of Brassica species

Seeds of 6 Brassica species (BF, LG, LM, DT, DE and DM) were sowed in 8x8x8.4 cm square pots filled with common potting mix soil (20 pots per species), distributed randomly in trays and placed in a controlled environment room (12 hr light, 22°C ± 2°C). Five days after germination, one plant per pot was left. Plants were irrigated with ¼ Hoagland solution every 2/3 days. Seven days post-germination, 3/4 of the plants (15 pots per species) were gradually irrigated with 25 mM of NaCl, 1/3 of them reaching 100 mM of NaCl and collected at 21 days old, 1/3 reaching 200 mM of NaCl and collected at 32 days old, and 1/3 reaching 300 mM of NaCl until maturation. The aerial tissue of each plant was dried for 48h at 60 °C and weighed to obtain the aerial biomass.

### Reciprocal transplant experiment

*Brassica fruticulosa* seeds from 3 coastal (BLA, ROS, TOS) and 3 inland (PAU, TOR, SOL) populations were sowed in uncultivated fields of the Marimurtra Botanic Garden at Blanes (JBB: 41.677320, 2.801198), a representative coastal environment, and of Les Planes d’Hostoles (LP: 42.064403, 2.544788), a representative inland environment. The same common garden design was reproduced at both sites, with 10 individuals per population planted into 20x20-cm squares. One leaf of three 30-days old plants per population was harvested and dried for ionomic analysis. Fitness was evaluated counting the number of siliques at maturity.

### Germination tests and hydroponic cultivation

*Brassica fruticulosa* seeds from 9 coastal (BEG, BLA, ESC, GAR, LAN, POR, ROS, SFG, TOS) and 9 inland populations (BRU, FUM, LOF, PAL, PAU, PiM, SCF, SOL, TOR) were first soaked in 30% bleach solution (30% bleach, 70% deionized (dd) water and 2 drops of tween) for 15 minutes, then washed with ddH_2_O 5 times, and finally suspended in 10^-5^ potassium nitrate solution. Seeds were stratified for 2 days at 4 °C to synchronize germination. 30 seeds from each population were placed in square plates with ½ MS medium (pH 6, Control) or ½ MS medium with 75 mM of NaCl (pH 6, Salt) to test the germination rate. Plates were placed in a growth chamber with 150 mmol/m^2^s of light intensity, 12h light/12h dark photoperiod, and 25 °C day/night temperature.

One-week old seedlings from the control plates were transferred to a hydroponic system filled with 0.5-strength Hoagland solution (pH 6, Control) in the same growth chamber. The hydroponic solution was changed every 2-3 days to maintain a constant concentration of nutrients in the solution and in half of the plants NaCl was increased gradually (50 mM NaCl every 2/3 days) until a final concentration of 150 mM of NaCl (pH 6) when plants were 21 days old. Ten days later, we harvested the plants and measured plant growth parameters (root length, biomass). Plants from 4 coastal (BLA, ESC, ROS, TOS) and 4 inland *B. fruticulosa* populations (PAU, SCF, SOL, TOR) were selected to measure osmotic potential, proline, and root and leaf ionome. Samples for RNA extraction were immersed in liquid nitrogen, homogenized to a fine powder, and stored at –80 degrees C.

### Soil and plant tissue ionomic analysis

To characterize the elemental composition of soil, analyses were performed on the 2-mm fraction samples following the extraction method described in Busoms et al. [1]. Plant tissue was dried for 2 days at 60 degrees C. Approximately 0.1g was weighed and used to performed open-air digestion in Pyrex tubes using 0.7 mL concentrated HNO_3_ at 110°C for 5h in a hot-block digestion system (SCI54-54-Well Hot Block, Environmental Express, Charleston, SC, US). Concentrations of the following elements (Li, B, Ca, K, Mg, Na, P, S, Mo, Cu, Fe, Mn, Co, Ni, Zn, Cr, As, Rb, Sr, Cd, and Pb) were determined by ICP-MS (Perkin Elmer Ink., ELAN 6000, MA, US) or ICP-OES (Thermo Jarrell-Ash, model 61E Polyscan, UK). Mean-standardized values (1 < value > 1) of elemental contents were used to represent the radar plots.

### Proline quantification

Proline concentration was determined colorimetrically from frozen tissue using a method adapted from Bates *et al.* [55]. Briefly, fresh plant material (50 mg) was homogenized with 3% 5-Sulfosalicylic acid and centrifuged at 10,000 rpm, 4°C for 10 min. Supernatant was collected and stored at -20°C. For the reaction mixture preparation, ninhydrin was dissolved in hot glacial acetic acid (1:24 w/v) and then carefully mixed with 6M orthophosphoric acid (3:2 v/v). In crystal tubes, glacial acetic acid, ninhydrin mix, and plant extract was mixed (1:1:1 v/v/v) and subjected to a hot water bath at 98°C for 60 min. Then, samples were cooled down in ice for 20 min and toluene was added and vortexed for 20 sec. Finally, the organic phase was extracted, and the absorbance was measured at 520 nm. Proline concentration was calculated using a calibration curve.

### Osmotic potential

Frozen tissue (0.2g) was placed into a 0,5ml Eppendorf and boiled in a water bath for 30 min. The resulting liquid was extracted, centrifugated 10 min at 15,000 rpm, and the supernatant was measured in a freezing-point depression osmometer (Osmomat 3000, Ganotech).

### Oxford Nanopore sequencing, Genome assembly and annotation

High molecular weight (HMW) DNA was isolated from one plant from Tossa de Mar (TOS), Catalonia, Spain. HMW DNA was extracted from 2.5 g of leaves that had been etiolated for 24 hours, according to the nuclear DNA isolation protocol published by Bernatzky and Tanksley [56], with some modifications. In brief, leaves were snap frozen in liquid nitrogen, ground to a fine powder, homogenized in extraction buffer (0.35 M Sorbitol, 100 mM Tris, 5 mM EDTA (pH 7.5) with 1 % β-mercaptoethanol added freshly before use) by vortexing for 30 s. The resulting slurry was filtered through Miracloth and centrifuged for 15 min at 720 g. The pellet was resuspended in extraction buffer containing 0.4 % Triton X-100 and centrifuged for 15 min at 720 g. This step was repeated a second time. The pellet, containing nuclei, was resuspended in extraction buffer containing 100 ug/ml of RNase and nuclei were lysed by mixing the suspension with CTAB Lysis Buffer (ITW Reagents; A4150) and adding 1/6 volume of 5 % N-laurylsarcosine. The lysis mix was incubated at 65 degrees C for 20 min, before the lysate was mixed with 2.2 volumes of chloroform/isoamylalchol (24/1) and centrifuged at 5000 g for 15 min. The aqueous phase was mixed with an equal amount of cold isopropanol to precipitate the DNA. The DNA was hooked out of the solution with a glass rod, washed by dipping in 80 % ethanol, and then resuspended in 100 ul of TE buffer (10 mM Tris-HCl, 1 mM EDTA; pH 8). Genomic DNA was left to resuspend overnight, then quantified using the Qubit Fluorometer and the Qubit DNA dsDNA BR Assay (ThermoFisher; Q32853). DNA (1 ug) was used as the input for an ONT ligation sequencing library preparation (ONT; SQK-LSK109). The library was quantified using the Qubit Fluorometer and the Qubit dsDNA HS Assay Kit (ThermoFisher; Q32854). 650 ng of library was sequenced over one PromethION flow cell (ONT; FLO-PRO002) and run on a PromethION Beta sequencer.

Fast5 sequences were basecalled using Guppy (version 6 high accuracy basecalling model dna_r9.4.1_450bps_hac.cfg; [57]) and the resulting fastq files were quality filtered by the base caller. Base called fastq files were assembled using Flye (version 2.9; [58]) at a depth of 38x, assuming a 557 megabase genome (Kmer-based genome size estimates were performed with FindGSE [20].) into a pseudohaploid primary assembly. Contigs were then polished to improve the single-base accuracy in a single round of polishing with Medaka (version1.5.0; [59]) using the ONT long reads, followed by a second round of polishing incorporating paired-end Illumina HiSeq data with Pilon (version 1.24) [22]. We purged uncollapsed duplicate haplotypes using PurgeDups [23]. We then used the BRAKER pipeline to conduct gene annotation on the genome assembly [26]. Evidence types included RNAseq data (generated as below) and protein data from related species.

### Contamination removal and assembly assessment

Removal of contamination from the assembly was performed in Blobtools [24], based on phylogenetic assignment, copy number, and GC content. The Uniprot database was downloaded (8 May 2022) and the assembly was blasted against this using (Diamond version 2.0.15) [60]. Gene space completeness was assessed using BUSCO version 5.2.2 [61] and the odb10 database for brassicales, employing default parameters.

### Short-read library preparation and sequencing

Genomic DNA was isolated from *B. fruticulosa* young leaves using the Wizard® Genomic DNA Isolation Kit. Quantification of DNA was performed using Quant iT dsDNA High Sensitivity Assay (Thermo Fisher; Q-33120) with a Fluoroskan Ascent Fluorometer (Thermo Fisher; 5200111). Illumina DNA sequencing libraries were constructed using Illumina DNA Prep library preparation kit (Illumina; 20018705) with IDT® for Illumina Unique Dual indexes, sets B and D Library preparation was performed using a Mosquito HV (SPT Labtech) liquid handling robot with 1/10^th^ volumes at all steps. A total of 9 – 48 ng of DNA was used as library input and 5 cycles of PCR were used for the library amplification step. Final libraries were normalised and pooled based on quantification performed by Quant iT dsDNA High Sensitivity Assay (Thermo Fisher; Q-33120) on a Fluoroskan Ascent Fluorometer (Thermo Fisher; 5200111). The resulting pools were size selected using 0.65X AMpure XP Beads (Beckman Coulter; A63882) to remove library fragments < 300 bp. Size selected pools were checked for size distribution on High Sensitivity D1000 ScreenTape (Agilent; 5067-5584) using an Agilent TapeStation 4200 and quantified using Qubit™ 1X dsDNA High Sensitivity (HS) Assay on a Qubit 4 Fluorometer. Final library pool quantification was performed using the KAPA Library Quantification Kit for Illumina (Roche; KK4824). The libraries were sequenced with Illumina NovaSeq 6000 at Novogene Europe, Cambridge, UK.

### Short-read processing and variant calling

Low quality reads and sequencing adapters were removed using Trimmomatic (PE -phred33 LEADING:10 TRAILING:10 SLIDINGWINDOW:4:15 MINLEN:50; [62]) and the surviving reads aligned to the *B. fruticulosa* refence genome using bwa-mem2 [63]. We removed duplicated reads using Picard tools (https://broadinstitute.github.io/picard/) and identified SNPs using GATK4 [64]. Filtering of the variant calls was based on the GATK’s best practice guidelines, and we included filters for mapping quality (MQ³ 40 and MQRankSum³ –12.5), variant confidence (QD³ 2), strand bias (FS < 60), read position bias (ReadPosRankSum³ –8), and genotype quality (GQ³ 10). We further removed SNPs with sequencing depth 1.6x the mean depth to avoid issues caused by paralogous mapping.

### Analysis of genetic structure and variation

All genetic structure and demographic analyses were run using fourfold degenerate sites identified with DEGENOTATE (https://github.com/harvardinformatics/degenotate). Performance was verified by running SNPeff to confirm the presence of only synonymous variants. For fastSTRUCTURE analysis we LD thinned in 5kb windows allowing no more than 20% missing data and minimum allele frequency of 2.5% to yield a dataset of 30,272 fourfold degenerate sites. fastSTRUCTURE was then run at K1 to K10 five times to clarify the extent of gene flow between populations. For PCA and splitsTree analysis of Nei’s genetic distances we thinned the dataset with a custom C script prune_ld [65] that thinned to *r*^2^ ≤ 0.1 in 100 SNP windows also allowing no more than 20% missing data and minimum allele frequency of 2.5% which yielded a dataset of 19,689 fourfold degenerate sites. We then conducted a principal component analysis (PCA) on the fourfold dataset ([MAF] > 0.05). Following Patterson et al., (2006), we estimated a covariance matrix representing the genetic relationships among each pair of individuals. For two individuals, *i* and *j*, covariance (*C*) was calculated as:

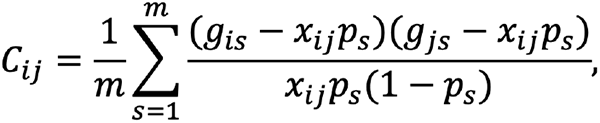

where *m* is the number of variable sites, *gis* is the genotype of individual *i* in site *s*, *x* is the average ploidy level of the two individuals, and *p* is the alternate allele frequency. We then conducted PCA on the matrix using the R function prcomp and extracted the first two axes of the rotated data for plotting.

### Sequence Processing and Allele Frequency Estimation

Likelihoods for the three possible genotypes in each biallelic site were then calculated from the BAM files in ANGSD 0.939 [66]. Nucleotides with base qualities lower than 20 and reads with mapping quality lower than 30 were discarded. Sites with coverage at fewer than 72 out of the 90 samples were also excluded. Analysis of genetic diversity and population structure was performed in ANGSD using the empirical Bayes (EB) method to calculate Tajima’s D [66], which approximates how far the population is from a mutation-drift balance [70]. Splitstree [71] was used to clarify the extent of gene flow between populations.

### Selection Scans

Allele frequencies for the selection scan were estimated directly from the site allele frequency (saf) likelihoods based on individual genotype likelihoods assuming HWE [72] in ANGSD. Fst was estimated using Hudson’s estimator as defined in [73], since it is independent of sample sizes even when Fst is not identical across populations. Selected sites were detected by scanning the contigs in 1kb non-overlapping windows for areas of localised extreme differentiation between the coastal and inland populations, which is a pattern indicative of directional selection [74]. The regions with the highest 1% values were identified and considered as candidate regions under selective sweeps. Fst distributions were plotted with ggplot2 package [75] using 1kb non-overlapping (step size 1000 bp) sliding windows. PCAdapt was run with default paramaters on the GATK-called, best practices filtered VCF [43].

### Transcriptome analysis

RNA from leaves and roots of salt-tolerant and salt-sensitive *B. fruticulosa* plants hydroponically cultivated under 0 mM or 150 mM NaCl was extracted using the Maxwell plant RNA kit (Promega Corporation, Madison, WI, USA) following the manufacturer’s instructions. Total RNA was quantified using a Qubit 2.0 Q32866 (Life Technologies, Carlsbad, CA, USA) and then used to prepare the complementary deoxyribonucleic acid (cDNA) library. cDNA library was composed by 48 samples (2 tissues x 2 treatments x 2 origins x 2 salinity phenotypes x 3 replications) and sequenced at the Illumina NovaSeq 6000 using standard procedures to generate 150 bp paired end reads (Novogene, Sacramento, CA, USA). Quality assessment of raw sequencing data was performed with FastQC software. Low-quality reads and sequencing adapters from raw data were removed with Trim Galore! software (https://github.com/FelixKrueger/TrimGalore)*. Rsubread* package was used to map the paired-end reads to the reference genome [76] and *samtools* [69] was used to sorting and indexing. HTSeq-counts was applied to count the overlap of reads with the gene models [77]. PCA of root and leaf data were performed on the whole transcript levels. The differential expression analysis was performed with *DESeq2* R Bioconductor package (1.39.2) [78]. Differential expressed genes (DEGs) were selected based on L2FC > |2|, p-adj < 0.05.

### Enrichment analysis

Gene Ontology (GO) terms were obtained with a customized ‘*B. fruticulosa* GO univers’ created with orthofinder [79] using orthologous genes of *A. thaliana* and *Raphanus sativus*. Fisher’s Exact Test was used to determine enrichment using elim algorithm in topGO packages of Bioconductor. STRING platform was used to visualize protein-protein interactions networks and perform enrichment analysis with the genes with A. thaliana orthologues. The transcription factors (TF) with significantly over-represented target number and the regulations between the TFs and the input genes were obtained using the ‘TF Enrichment’ online tool of the Plant Transcriptional Regulatory Map (https://plantregmap.gao-lab.org/tf_enrichment.php).

### Environmental association analysis

To identify candidate genes associated with particular soil ionome characteristics, we performed environmental association analysis (EAA) using Latent Factor Mixed Models (LFMMs)—LFMM 2 (https://bcm-uga.github.io/lfmm/; [42]) using the soil and leaf ionome as phenotype. We tested the association of allele frequencies at each SNP for each individual with associated soil concentration of elements differentiating saline and non-saline soils. We retained 1,080,632 SNPs without missing data and MAF > 0.05 as an input for the LFMM analysis. LFMM accounts for a discrete number of ancestral population groups as latent factors. We used 3 latent factors reflecting the number of genetics clusters in our dataset. To identify SNPs significantly associated with soil variables, we have used a p-value threshold of 1×10^-5^. Finally, we annotated the candidate SNPs to genes, termed ‘LFMM candidates’ (at least one significantly associated SNP per candidate gene).

### Climatic data

Monthly precipitation (mm), reference evapotranspiration (ET_0_, mm), and maximum, minimum and mean temperature (°C) from 2010 to 2023 was obtained from the ‘Servei Metereologic de Catalunya’ (https://ruralcat.gencat.cat/web/guest/agrometeo) selecting Roses and Cabanes stations for the north region and Malgrat de Mar and Viladrau stations for the central region. ET_0_, or theoretical evapotranspiration in a soil covered with a uniform 10 cm high turf cover, was calculated according to the Penman-Monteith methodology [80].

### Statistical analysis

One-way or multivariate analysis of variance (ANOVA/MANOVA) were used to test for significant differences (p < 0.05) between means of data with respect to growth, physiological parameters, ionome, fitness, gene expression, and climatic data. To test for correlations between two variables, a bivariate fit was applied. To perform multiple comparisons of group means, Tukey’s HSD test was conducted. A permutation test was performed to determine whether the number of overlaps between GO categories observed was likely to occur by chance. For each condition (Fst, EAA, PCAd and RNA) a list of random genes the same size as the real candidate gene list was generated using all the genes in the genome. GO analyses were then performed on these random lists and the number of overlapping GO categories was determined. This was performed 10,000 times for both the north and central candidate gens lists. P-values were calculated as X/10,000 where X is the number of times there was an equal or greater number of overlapping GO terms in the randomized test versus the real data (Dataset S11). Statistical analyses and plots were performed using SAS Software JMP (v.16.0) or R (version 4.2).

## Acknowledgements

This work was supported by the European Research Council (ERC) under the European Union’s Horizon 2020 research and innovation programme [grant number ERC-StG 679056 HOTSPOT], via a grant to L.Y.; the Biotechnology and Biological Sciences Research Council [grant number BB/P013511/1], via a grant to the John Innes Centre, and via a grant BB/R017174/1 to L.Y.; and the Spanish Ministry of Science and Innovation Project PID2019-10400RB-I00. We acknowledge support also from The University of Nottingham, in particular through the Future Food Beacon of Excellence. Computational resources were provided by the e-INFRA CZ project (ID:90254), supported by the Ministry of Education, Youth and Sports of the Czech Republic. We thank Michael Giolai for the help and support during field collections and Andrew MacColl for helpful comments on the manuscript.

## Author Contributions

LY and SBu conceived the study. SBu, LY, AdS, EC, GE, RA, SBr and MW performed data analyses. SB, GE, ABG performed laboratory experiments. SBu, CP and ABG performed collections. LY and SBu wrote the manuscript with input from all authors. All authors edited and approved the final manuscript.

## Data availability

PromethION, Illumina sequencing, and RNAseq raw data generated in this study have been deposited in the European Nucleotide Archive under the project accession PRJEB74663. The genome assembly and annotations generated in this study have been deposited on Zenodo [https://zenodo.org].

