## Supplementary material for "Local cryptic diversity in salinity adaptation mechanisms in a wild outcrossing *Brassica*": SI figures

### Supplementary Figures

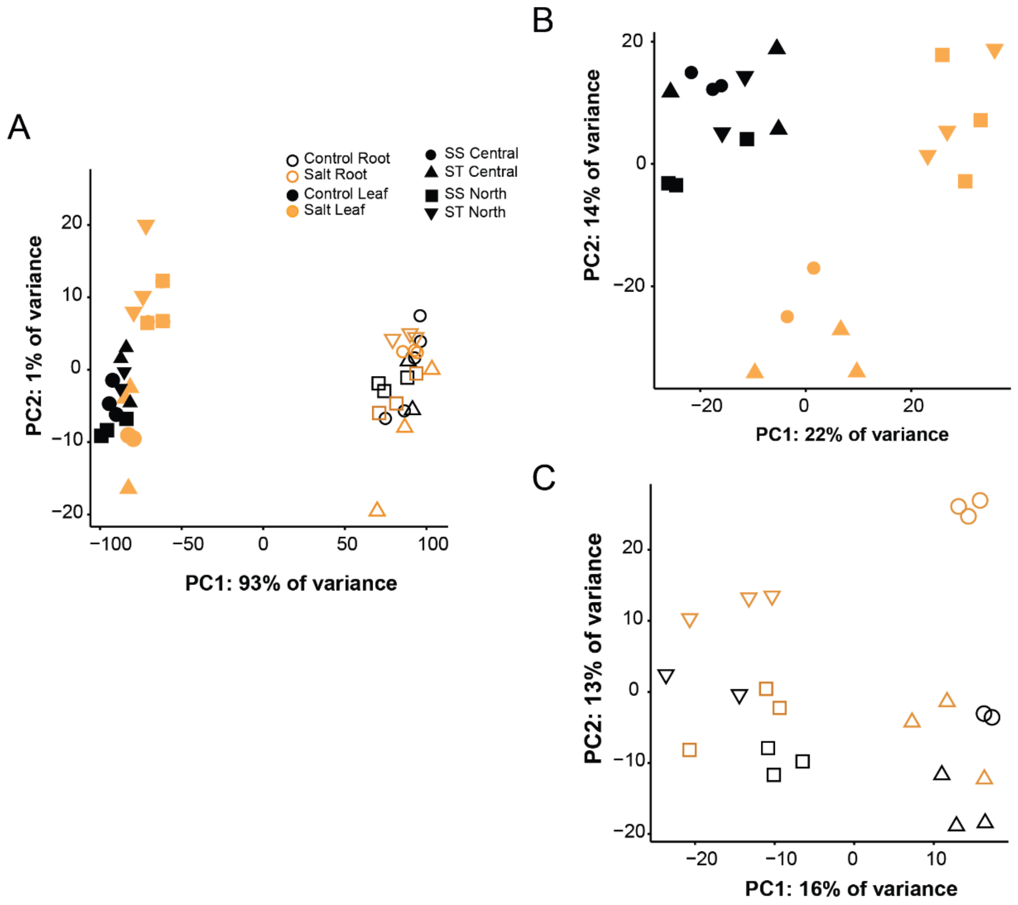


**Figure S1.** PCA of **(A)** all *Brassica fruticulosa* transcripts, **(B)** of leaf transcriptome profiles, and **(C)** root transcriptome profiles of 12 salt-tolerant (triangles up/down) and 12 salt-sensitives (circle/square) *B. fruticulosa* individuals from north and central metapopulations treated with 0 mM NaCl (control, black) or 150 mM NaCl (salt, orange) for 10 days.

A

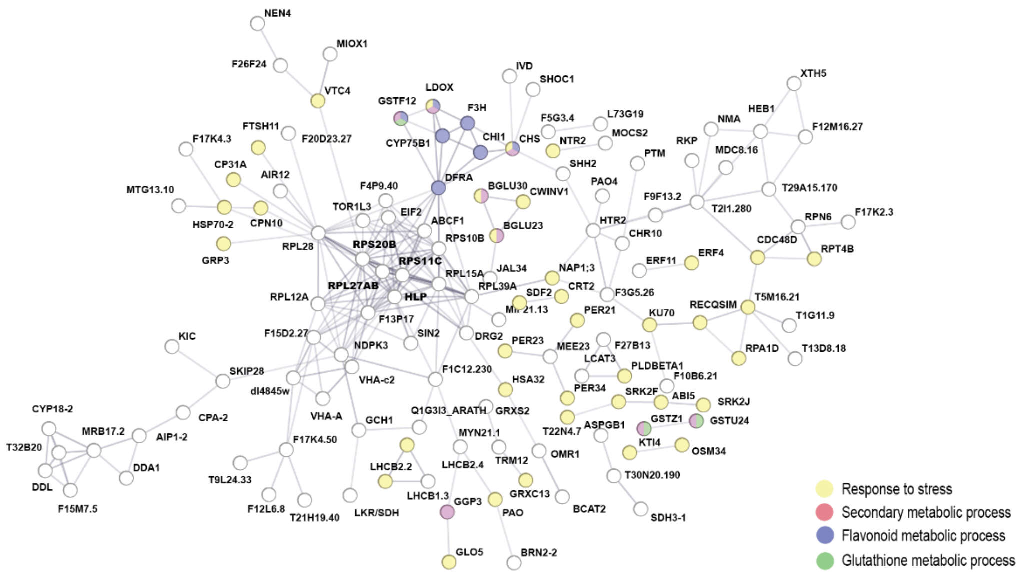


B

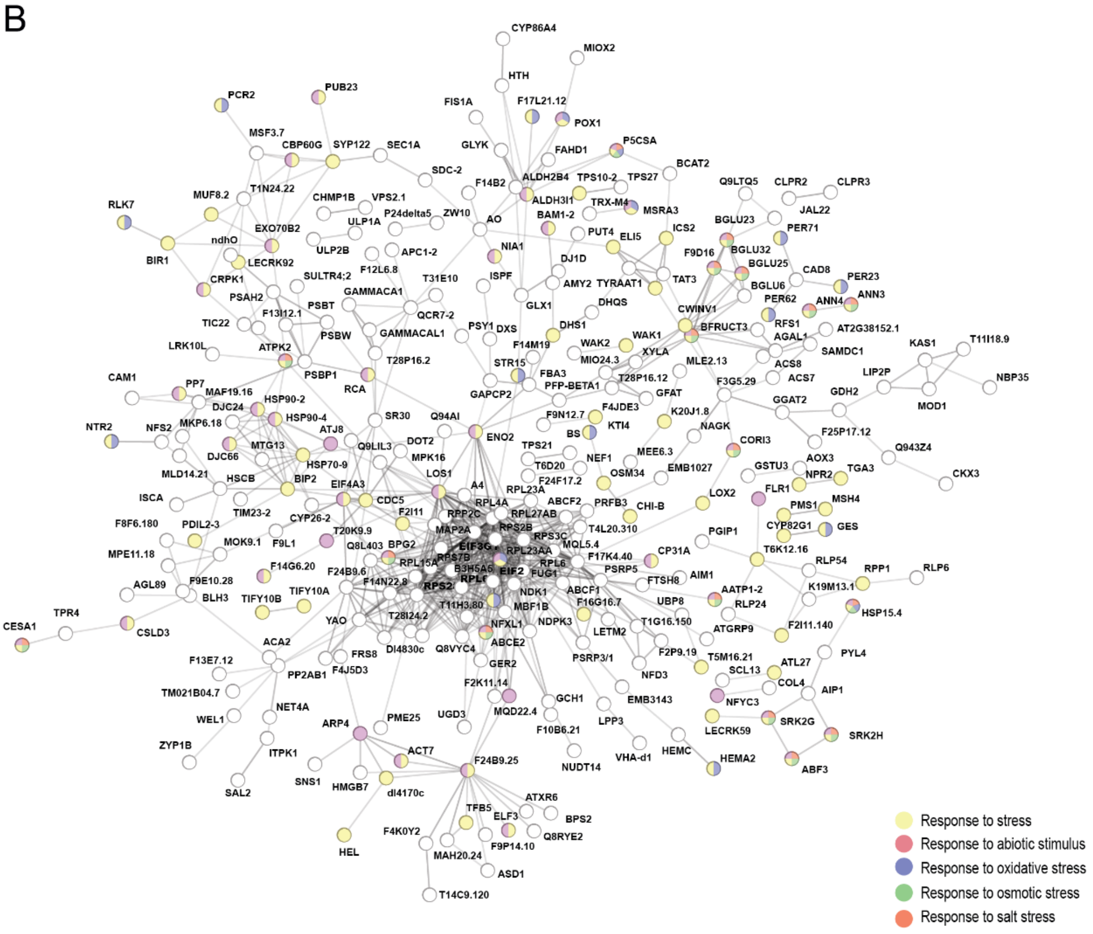


**Figure S2.** STRING networks of DEGs from the pairwise comparations salt tolerant *vs* salt sensitive populations under salt stress of **(A)** north populations **(B)** central populations. Each sphere corresponds to one gene and nodes represent protein-protein interactions Only connected proteins are represented and proteins from relevant enriched GO terms are coloured.

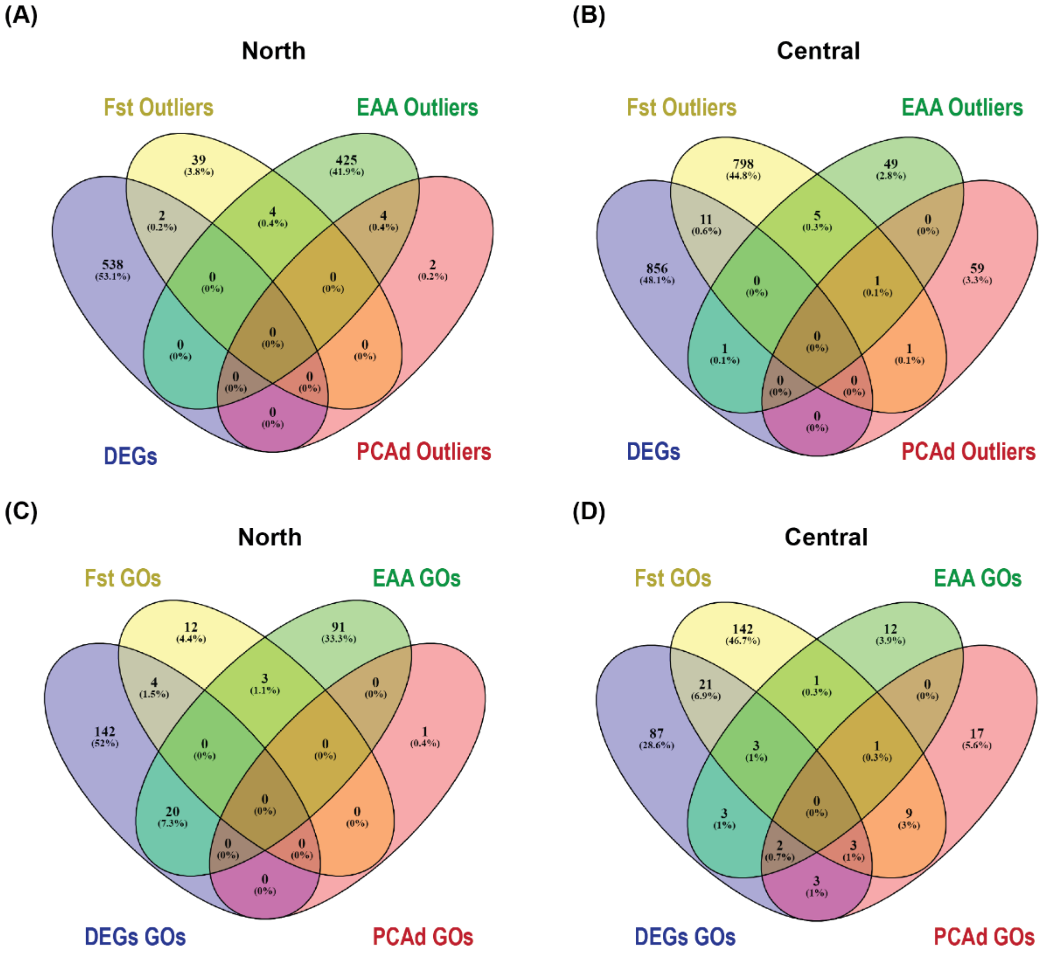


(E)

**North**

**Common Terms RNA and EAA outliers (**)**
carbohydrate metabolic process
cellular response to organic substance
cytokinin-activated signaling pathway
defense response to other organism
developmental process
hormone-mediated signaling pathway
immune response
inorganic anion transport
macromolecule metabolic process
regulation of gene expression
response to endoplasmic reticulum stress
response to lipid
response to nematode
*response to osmotic stress*
response to wounding

**Common Terms RNA and Fst outliers (*)**
negative regulation of metabolic process
recognition of pollen
response to ethylene

**Common Terms Fst and EAA outliers (*)**
cellular developmental process
cellular metabolic process
cellular process

(F)

**Central**

**Common Terms RNA and Fst outliers (**)**
carbohydrate derivative biosynthetic process
cell wall biogenesis
cellular response to lipid
cellular response to organic substance
cellular response to oxygen-containing compound
defense response to other organism
developmental growth
DNA integration
embryo development ending in seed dormancy
endosome transport via multivesicular body sorting pathway
hormone-mediated signaling pathway
negative regulation of metabolic process
organonitrogen compound biosynthetic process
photoperiodism, flowering
protein folding
regulation of DNA-templated transcription
regulation of pH
response to abscisic acid
response to light stimulus
response to lipid
*response to salt stress*
*response to stress*
*response to water*
*response to water deprivation*
small molecule biosynthetic process
stomatal movement

**Common Terms RNA and EAA outliers (**)**
auxin-activated signaling pathway
jasmonic acid mediated signaling pathway
seed development
stamen development

**Common Terms EAA and PCAdapt outliers (*)**
jasmonic acid mediated signaling pathway
cellular process

**Common Terms Fst and EAA outliers**
cell growth
cellular process

**Common Terms fst and PCAdapt ouliers (**)**
defense response to bacterium
response to bacterium
cell wall organization or biogenesis
protein metabolic process
nitrogen compound metabolic process
response to organic substance
primary metabolic process
cellular metabolic process

**Common Terms RNA and PCAdapt ouliers (*)**
defense response to fungus
glutathione metabolic process
macromolecule metabolic process
vesicle-mediated transport

**Common Terms RNA, Fst and EAA outliers (**)**
cellular response to organic substance
embryo development ending in seed dormancy
*response to water deprivation*

**Common Terms RNA, Fst and PCAdapt outliers (**)**
defense response to other organism
negative regulation of metabolic process
*response to abscisic acid*
response to stress

**Common Terms RNA, EAA and PCAdadapt outliers (**)**
jasmonic acid mediated signaling pathway

**Common Terms Fst, EAA and PCAdapt outliers (**)**
cellular process

**Figure S3.** Candidate genes overlaps of *B. fruticulosa* north (**A**) and central (**B**) genomic (Fst, EAA, and PCAdapt outliers) and transcriptomic (DEGs) analysis. Overlap of the enriched GO terms of the candidate genes from north (**C, E**) and central (**D, F**) genomic and transcriptomic analysis. Asterisks indicate significant contrasts (Permutation test; * < 0.05 threshold, ** < 0.01 threshold).

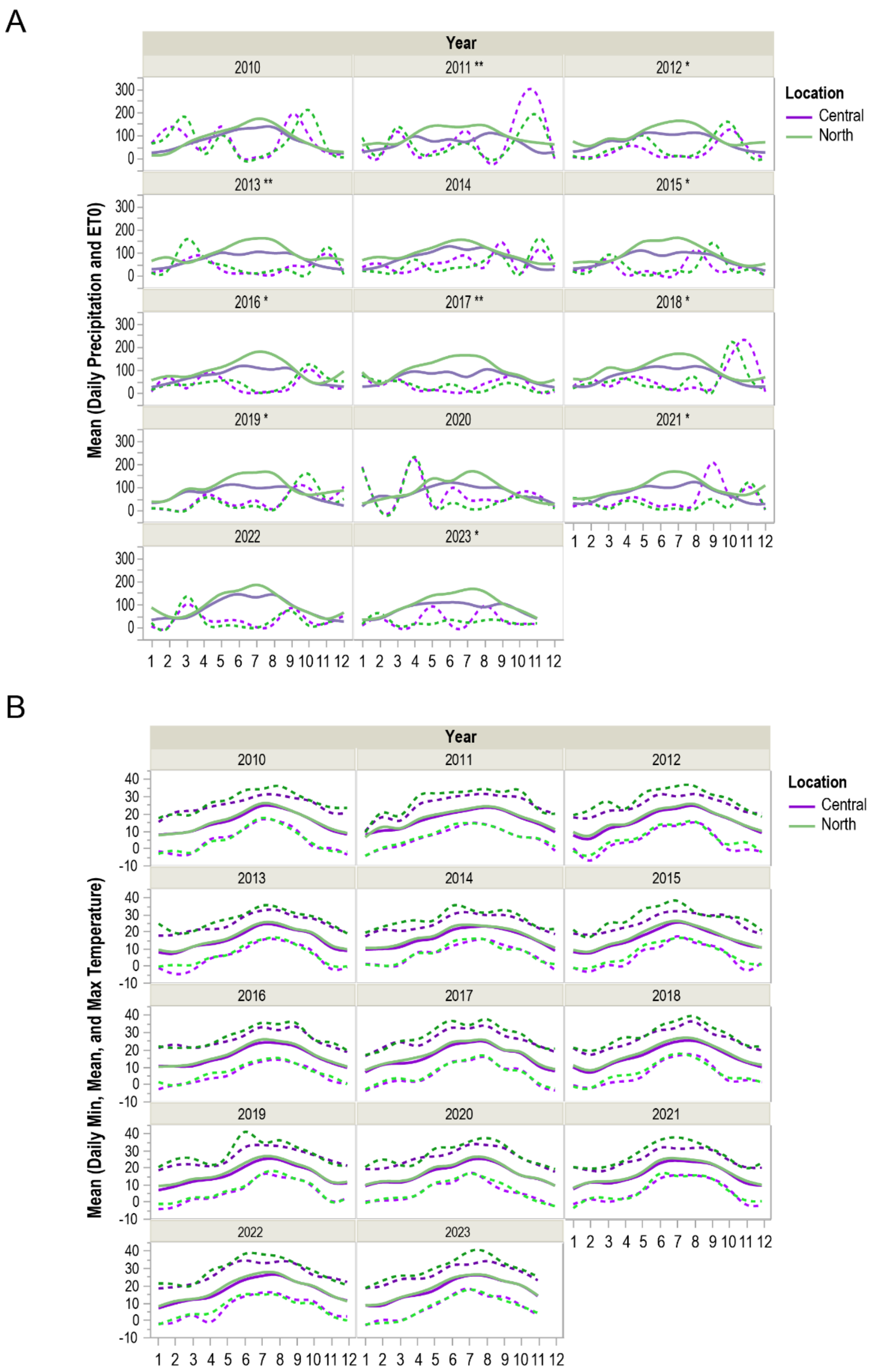


**Figure S4. (A)** Monthly mean precipitation and **(B)** monthly mean minimum, mean and maximum daily temperature of 14 years (2010-2023) recorded in Roses (north, green line) and in Malgrat de Mar (central, purple line) stations.
